# Axonal ER tubules regulate local translation via P180/RRBP1-mediated ribosome interactions

**DOI:** 10.1101/2022.11.30.518484

**Authors:** Max Koppers, Nazmiye Özkan, Ha H. Nguyen, Daphne Jurriens, Janine McCaughey, Riccardo Stucchi, Maarten Altelaar, Lukas C. Kapitein, Casper C. Hoogenraad, Ginny G. Farias

**Author notes:** Correspondence (G.G.F).

## Abstract

Local mRNA translation in axons is critical for the spatial and temporal regulation of the axonal proteome. A wide variety of mRNAs are localized and translated in axons, however how protein synthesis is regulated at specific subcellular sites in axons remains unclear. Here, we establish that the axonal endoplasmic reticulum (ER) supports axonal translation. Axonal ER tubule disruption impairs local translation and ribosome distribution. Using nanoscale resolution imaging, we find that ribosomes make frequent contacts with ER tubules in the axon in a translation-dependent manner and are influenced by specific extrinsic cues. We identify P180/RRBP1 as an axonally distributed ribosome receptor that regulates local translation in an mRNA-dependent manner. Our results establish an important role for the axonal ER in localizing mRNA translation and in dynamically regulating the axonal proteome in response to neuronal stimuli.

## INTRODUCTION

The complex and polarized nature of neurons requires precise regulation of the local proteome to maintain neuron development and function. mRNA localization and translation in dendrites and axons are essential for tight spatiotemporal control of the local proteome in response to local demands.^1^ Recent studies have shown that axons contain and translate a diverse set of mRNAs, often in response to extracellular signals, which is important for many neuronal processes including axon guidance, synapse/branch formation, synaptic function and survival.^2–4^ Despite recent progress on axonal mRNA transport and translation mechanisms^5^, we still poorly understand where axonal protein synthesis takes place at a subcellular level and how this localization is achieved and regulated to fulfill local demands.

An unexplored player in local translation is the axonal endoplasmic reticulum (ER). Axons are devoid of rough ER or ER sheets and contain only ER tubules, which are presumed not to be involved in translation.^6,7^ However, studies in non-neuronal cells have shown that ribosomes, which bind ER sheets for translation of transmembrane, secretory, and a portion of cytosolic proteins^8^, can also bind ER tubules.^9,10^ In addition, some ER proteins involved in translation, which normally reside on ER sheets, have been detected in axons.^11,12^ However, whether axonal ER tubules have a functional role in local translation remains unknown.

Here, we establish a clear role for axonal ER tubules in the regulation of local mRNA translation. We demonstrate that the axonal ER interacts with ribosomes at the nanoscale resolution, and these contacts are sites for local translation in the axon. We reveal an extrinsic cue-specific regulation of these ER – ribosome contacts, suggesting an important role of the axonal ER in mediating stimuli-dependent translational regulation. We identify the integral ER protein P180 (also named RRBP1) as an axonally distributed ribosome/mRNA receptor, which facilitates ER-associated local translation. Mechanistically, P180 binds ribosomes in a mRNA-dependent manner through two binding sites, which together are sufficient to bind ribosomes and are required for efficient axonal ER – ribosome interactions. Together, our data indicate that the axonal ER, facilitated by P180, plays a critical role in regulating the subcellular localization of axonal mRNA translation.

## RESULTS

### The axonal ER regulates local mRNA translation and ribosome levels in axons

The ER appears continuous throughout the entire neuron consisting of two distinct membrane organizations. ER sheets, decorated with polysomes, are distributed in the somatodendritic domain and excluded from the axonal domain, while ER tubules are present in both domains.^6,7^ Axonal ER tubules are distinct from somatodendritic ER tubules; just a few long and narrow tubules without much complexity are observed in the axon.^13^

To test the idea that the axonal ER is involved in local translation, we first examined whether disruption of ER tubule formation affects axonal mRNA translation in primary cultures of rat hippocampal neurons. Knockdown (KD) of the ER tubule-shaping proteins RTN4 and DP1 causes a drastic reduction of the axonal ER, which is retracted and inter-converted to ER sheets in the somatodendritic domain.^12,14^ Thus, we performed short-hairpin RNA (shRNA)-mediated KD of RTN4 and DP1 and labelled overall axonal translation using the puromycilation assay^15^ at DIV7. After a short 10-minute puromycin incubation, newly synthesized proteins were labeled with an anti-puromycin antibody and quantified in a distal part of the axon (**Figure 1A; Figure S1A**). Compared to pSuper control neurons, RTN4/DP1 knockdown neurons showed a ±29% reduction in axonal puromycin labeling (**Figures 1B and 1C**).

**Figure 1.**
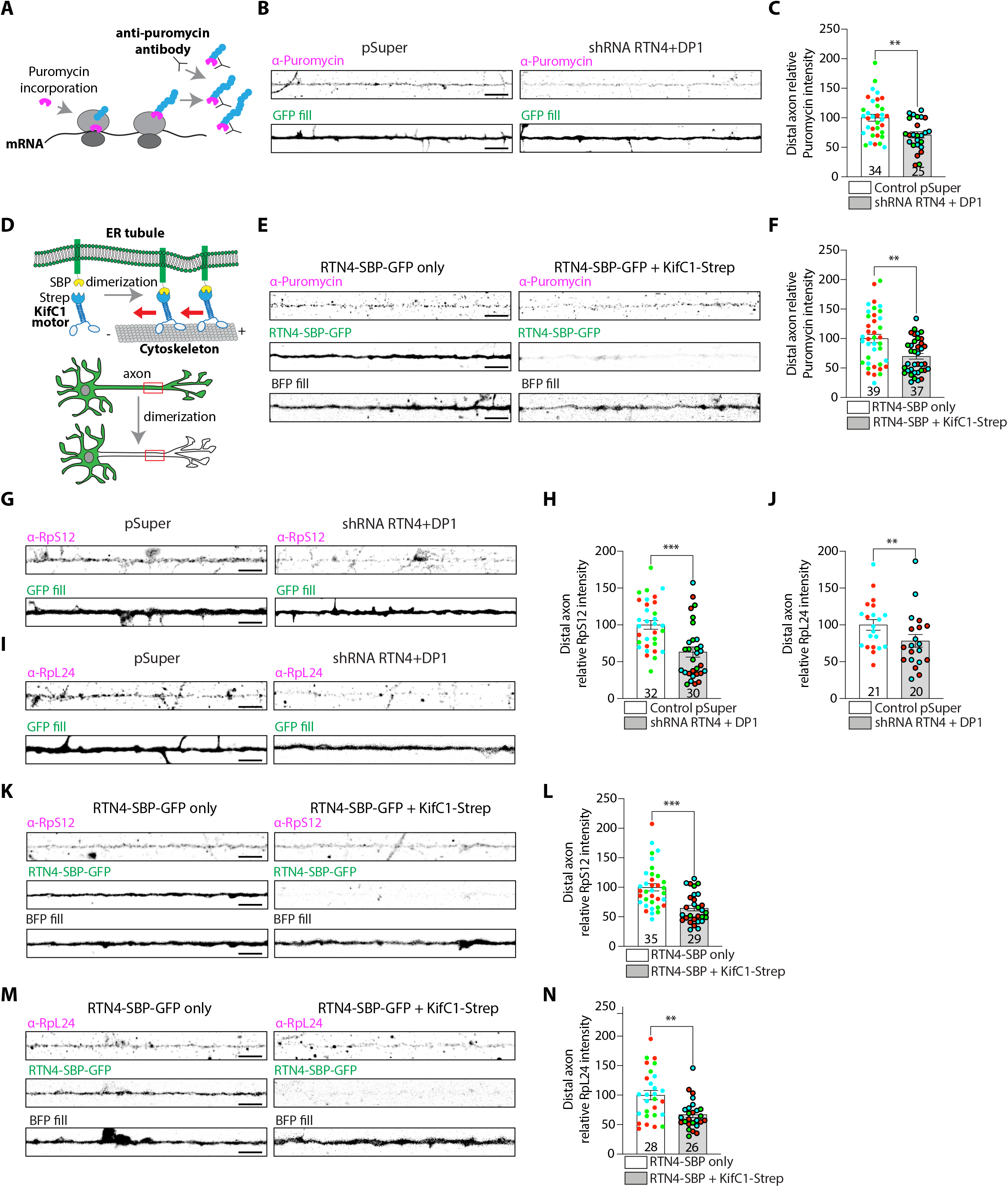
The axonal ER regulates local translation and ribosome levels. (A-C) Schematic showing puromycin incorporation into newly synthesized proteins (A). Representative images of puromycilated peptides in distal axons of DIV7 neurons transfected with a fill and a pSuper plasmid (control) or pSuper plasmids containing shRNA targeting RTN4 and DP1 (B). Quantification of puromycin intensity (C). (D-F) Schematic showing Streptavidin(Strep)-SBP heterodimerization system using SBP-RTN4 and Strep-KifC1 motor to relocate axonal ER into the soma (D). Representative images of puromycilated peptides in distal axons of DIV7 neurons expressing a fill and RTN4-SBP-GFP in absence or presence of Strep-KifC1 (E). Quantification of puromycin intensity (F). (G-J) Representative images of the distribution of endogenous ribosomal proteins RpS12 (G) and RpL24 (I) in the distal axon of neurons transfected as in (B). Quantification of RpS12 (H) and RpL24 (J) intensities from conditions as in (G) and (I), respectively. (K-N) Representative images of the distribution of ribosomal proteins RpS12 (K) and RpL24 (M) in distal axons of neurons transfected as in (E). Quantification of RpS12 (L) and RpL24 (N) intensities from conditions as in (K) and (M) respectively. Individual data points each represent a neuron and each color represents an independent experiment. Data are presented as mean values ± SEM in (C, F, H, J, L and N). *p < 0.05, **p < 0.01, ***p<0.001 comparing conditions to control with Mann-Whitney tests. Scale bars represent 5 μm.

Somatodendritic ER tubules are also disrupted by RTN4/DP1 KD.^14^ To study the local role of ER tubules in the axon, we used a heterodimerization system to selectively remove ER tubules from the axon.^12^ In this system, a streptavidin module is coupled to the minus-end directed motor KifC1 (Strep-KifC1) and a streptavidin binding protein (SBP) to GFP-tagged RTN4 (RTN4-SBP-GFP), which triggers their interaction and results in sustained removal of ER tubules from axons (**Figure 1D; Figure S1B**).^12,14^ We found that puromycin intensity in the distal axon decreased by ±30% after selective ER removal, similar to RTN4/DP1 KD (**Figures 1E and 1F**). Axonal ER removal did not affect the amount of newly synthesized proteins in the soma (**Figure S1C**). These results show that axonal ER tubules contribute to local translation in the axon.

If the axonal ER is directly involved in translation, one would expect that disruption of axonal ER tubules would also affect ER-bound ribosomes and thus the number of ribosomes present in the axon. We therefore studied the distribution of ribosomes along the axon by labelling the endogenous proteins RpS12 (eS12) and RpL24 (eL24) which are mainly present in the small and large ribosomal subunit, respectively.^16^ We first disrupted ER tubules by RTN4/DP1 KD and quantified ribosomal proteins in the distal axon. Compared to pSuper control, RTN4/DP1 KD resulted in ±37% and ±22% fewer ribosomes in the axon based on RpS12 and RpL24 staining, respectively (**Figures 1G-J**). We then specifically removed ER tubules from the axon using the Strep-SBP heterodimerization system. Similar to RTN4/DP1 KD, the selective ER removal from the axon resulted in a significant decrease of both RpS12 (±35% decrease) and RpL24 in the axon (±33% decrease) (**Figures 1K-N**). Together, disruption of axonal ER tubules impairs translation and ribosome distribution along the axon, suggesting a direct interaction between axonal ER tubules and ribosomes.

### Super-resolution imaging reveals the axonal ER frequently contacts ribosomes in a translation-dependent manner

We next examined whether the role of axonal ER tubules in local translation is due to a direct contact of ER tubules with ribosomes. We set out to visualize the possible association between the axonal ER and ribosomes at nanoscale resolution using three different super-resolution microscopy techniques.

To this end, we expressed the general ER marker Sec61β (GFP-tagged), present along the axonal ER, and we co-stained for the endogenous ribosomal protein RpS12. We first used stimulated emission depletion (STED) microscopy, which allowed us to resolve axonal ER tubules, consistent with what has been observed with FIB-SEM and Cryo-ET^6,17^, and generated a punctate signal for ribosomes (**Figure 2A**). Imaging of axonal segments and quantification revealed that ∼40-50% of ribosomes are in close proximity to the axonal ER (**Figure 2B and 2C**). These contacts are not random since horizontally flipping one channel resulted in a significantly lower overlap (**Figure 2C**). In addition to STED microscopy, we utilized two other super-resolution techniques, ten-fold robust expansion (TREx) microscopy^18^ and dual-color single-molecule localization microscopy (SMLM) using probability-based fluorophore classification.^19^ TREx and SMLM revealed a similarly high and specific close association of ribosomes with the axonal ER (**Figure 2D-I**). Thus, three different super-resolution imaging techniques show, at nanoscale resolution, consistent results indicating that a large portion of axonal ribosomes are associated with the ER in axons.

**Figure 2.**
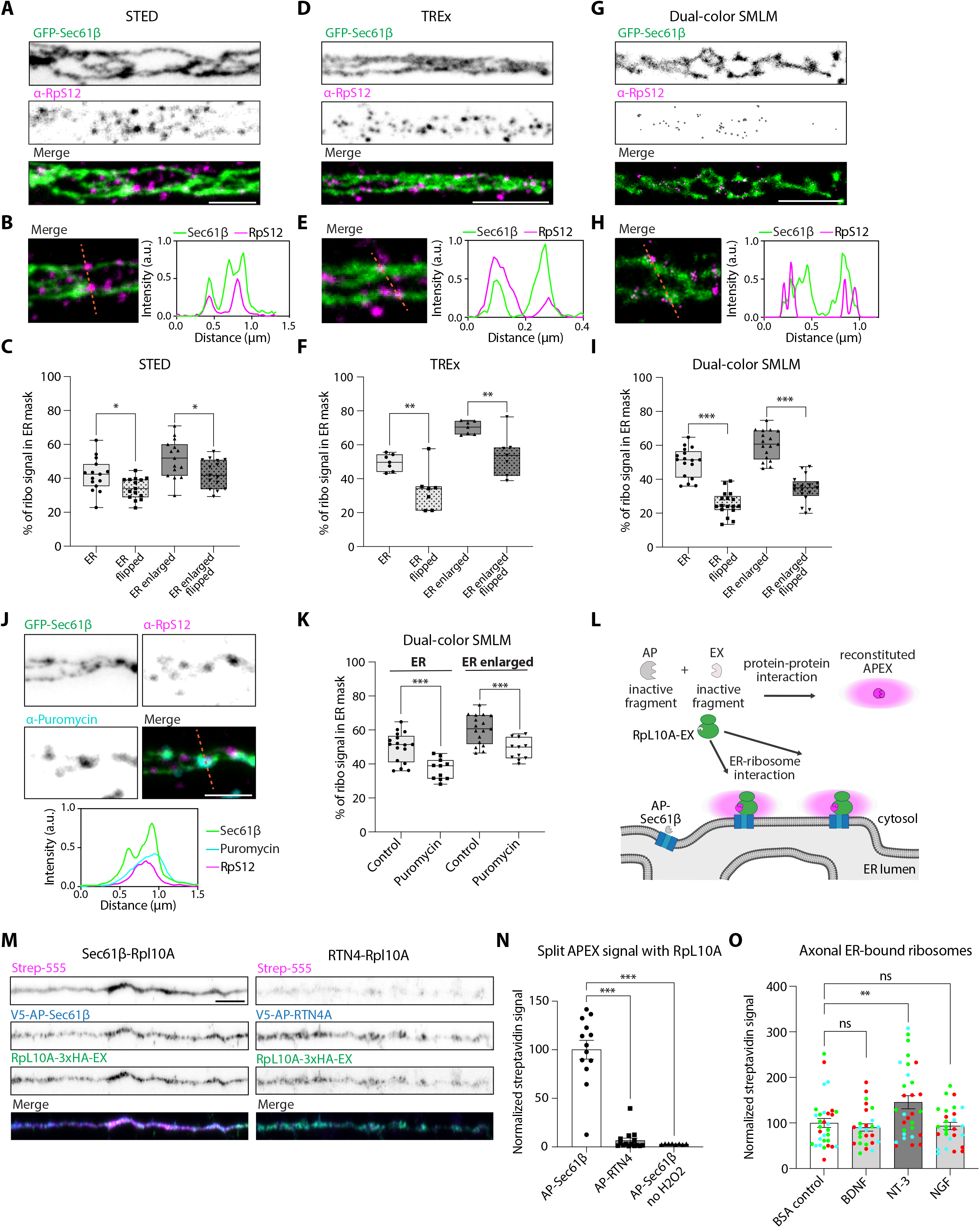
Nanoscale-resolved translation-dependent axonal ER – ribosome contacts. (A, D, G) Representative STED (A), TREx (D) and dual-color SMLM (G) images of the ER and ribosomes in axons from neurons expressing GFP-Sec61β and stained for RpS12. (B, E, H) Magnifications and intensity profile lines from merged images for each microscopy method. (C, F, I) Quantification of RpS12 intensity in ER mask, enlarged ER mask, and one-color flipped images, for each microscopy method. (J) Representative STED images and intensity profile line for an axon segment of a neuron transfected as in (A) and co-labeled for puromycilated peptides. (K) Quantification of RpS12 intensity in ER mask and enlarged ER mask with or without high puromycin treatment using dual-color SMLM. (L-N) Schematic representation of split APEX system used to detect ER-ribosome contacts. When the ER protein Sec61β fused to AP module and the ribosomal protein RpL10A fused to EX module interact with each other, APEX is reconstituted and contact sites can be visualized as a biotinylation radius around the interactions (L). Representative images of split APEX assay in distal axons from neurons expressing RpL10A-3xHA-EX and V5-AP-Sec61β (left), or V5-AP-RTN4A as a negative control (right). Expression of constructs are visualized with V5 and HA antibodies, and biotinylation is detected with conjugated Strep-555 (M). Quantification of Strep signal in distal axons from neurons as in (M), and without H2O2 as a negative control for the biotinylation reaction (N). (O) Quantification of axonal ER-bound ribosomes using split APEX assay with RpL10A-3xHA-EX and V5-AP-Sec61β, in neurons stimulated for 30-minutes with BSA (control), BDNF, NT-3 or NGF. Individual data points each represent a neuron in (C, F, I, K, N and O). Boxplots show 25/75-percentiles, the median, and whiskers represent min to max in (C, F, I and K). Data are presented as mean values ± SEM in (N, O). ns = not significant, *p < 0.05, **p < 0.01, ***p<0.001 comparing conditions to control using unpaired t-tests or ordinary one-way ANOVA tests. Scale bars represent 1μm (A, D, G, J) and 5μm (M).

Next, using STED, we visualized the axonal ER, ribosomes and sites of translation using the puromycilation assay. Interestingly, we observed various instances where ribosomes attached to the axonal ER were also positive for newly synthesized proteins (**Figure 2J**). This suggests that axonal ER – ribosome interactions are sites for local translation. To further confirm this, we treated neurons with a high concentration of puromycin, which causes ribosome disassembly and release of small subunits from the ER.^20,21^ Visualization and quantification of dual-color SMLM revealed a significant decrease of RpS12 associated to the axonal ER after high puromycin treatment (**Figure 2K**). This is consistent with bona fide contacts between the axonal ER and ribosomes and indicates that the axonal ER is a subcellular site for local translation.

To provide further evidence for a direct interaction, we utilized the split APEX system^22^, previously adapted to visualize organelle-organelle contacts in neurons.^14^ Theoretically, ER-bound translating ribosomes bind the ER translocon and we therefore attached an inactive AP-fragment to the translocon subunit Sec61β and an EX-fragment to the ribosomal protein RpL10A (uL1) (**Figure 2L**). These two fragments only reconstitute and enable biotinylation of nearby proteins when there is a molecular interaction between the ER and ribosomes. We first validated this system in neuronal soma where ER-bound ribosomes should be present. Quantification of somatic streptavidin labeling confirmed a specific interaction between Sec61β and RpL10A (**Figure S2A and S2B**). We then imaged axonal segments and quantified streptavidin labeling. Although the intensity of streptavidin labeling was expectedly lower than in the soma, we observed clear biotinylation with AP-Sec61β and RpL10A-EX in the axon, which was not seen with AP-RTN4 and RpL10A-EX (**Figure 2M**). Quantification of axonal segments confirmed the specific contact between the ER and ribosomes in the axon (**Figure 2N**).

Next, we explored whether ribosome contacts with axonal ER tubules are regulated by neuronal stimuli. Extrinsic signals are well known to trigger and enhance axonal mRNA translation.^3,23^ We shortly stimulated neurons with brain-derived neurotrophic factor (BDNF), neurotropin-3 (NT-3) or nerve-growth factor (NGF) and quantified ER – ribosome contacts using the split APEX assay. This revealed a cue-specific increase in axonal ER – ribosome interactions induced by NT-3, but not by BDNF or NGF (**Figure 2O**). Altogether, these results indicate that the axonal ER directly interacts with ribosomes in a translation-dependent manner, and this interaction is regulated by specific neuronal stimuli. Therefore, we propose that the axonal ER serves as a platform for local protein synthesis in the axon.

### P180 is an axonal ribosome receptor that facilitates axonal ER – ribosome interactions and local translation

We next sought to gain more insight into the proteins regulating axonal ER – ribosome interactions. Ribosomes can engage with the ER in signal recognition particle (SRP)-dependent and -independent manners.^24,25^ Besides the translocon, various ER-resident ribosome receptors have been proposed to play a role in ER-ribosome interactions and we therefore investigated the presence and/or enrichment of these proteins in the axonal compartment. We analyzed the subcellular distribution of four of these ER membrane proteins (P180, Leucine-rich repeat-containing protein 59 (LRRC59), Ribophorin 1 (RPN1) and Kinectin1 (KTN1); **Figure S3A**) as well as the translocon subunit Sec61β by quantifying the polarity index (PI; see Methods). Consistent with our previous study^12^ and our super-resolution microscopy data, Sec61β had a mostly unpolarized distribution (PI ∼ 0.28) and was present in both dendrites and the axon (**Figure 3A and 3B**). P180 was present in the soma but enriched in the axon and nearly absent from dendrites, consistent with previous findings (PI ∼ -0.4; **Figure 3A and 3B**).^12^ By contrast, LRRC59, RPN1 and KTN1 were strongly polarized towards the somatodendritic domain and were nearly absent from the axon (PI ∼0.7-0.8; **Figure 3A and 3B**). This suggests that, in addition to Sec61β, P180 may play a role in the interaction between the axonal ER and ribosomes. We therefore decided to explore the role of P180 in these interactions and in axonal translation.

**Figure 3.**
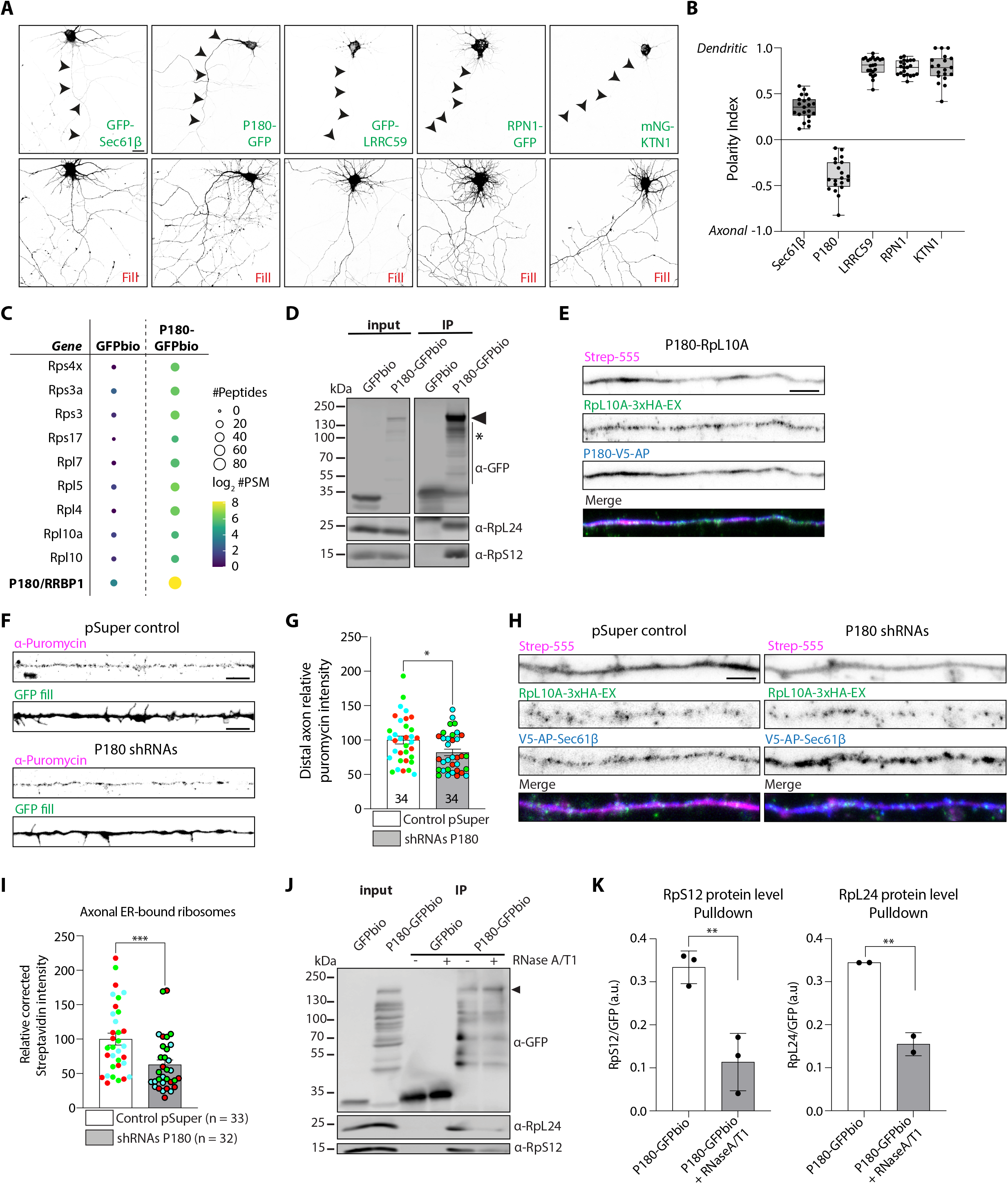
ER receptor P180 is enriched in axons and regulates local translation. (A-B) Representative images of neurons expressing a fill and the ER resident proteins GFP-Sec61β, P180-GFP, GFP-LRRC59, RPN1-GFP or mNG-KTN1 (A), and respective quantification of polarity index (B). (C) Scaled representation of proteins identified with mass spectrometry after pulldown of GFPbio or P180-GFPbio from adult rat brain extracts. The size and color of each dot reflect the number of PSMs or peptides identified as indicated in the legend. (D) Western blot validation for ribosomal protein interactions with P180-GFPbio after streptavidin pulldown. The presence of the coiled-coil domain in P180 causes protein instability resulting in a banded pattern, as previously described.^29^ (E) Representative images of split APEX assay for P180-V5-AP and RpL10A-3xHA-EX in the axon. Expression of constructs are visualized with V5 and HA antibodies, and interactions visualized with conjugated Strep-555. (F-G) Representative images of puromycilated peptides in distal axons from neurons expressing a fill plus control pSuper plasmid or shRNAs targeting P180 (F). Quantification in (G). (H-I) Representative images of split APEX in distal axons for V5-AP-Sec61β and RpL10A-3xHA-EX, in the presence of control pSuper plasmid or shRNAs targeting P180 (H). Protein expression and interaction visualized as in (E). Quantification in (I). (J-K) Western blot analysis of ribosomal proteins after GFPbio or P180-GFPbio pulldown with and without RNAseA/T1 treatment. Quantification in (K). Individual data points each represent a neuron (B, G and I), or an independent experiment (K). Data are presented as mean values in ± SEM. *p < 0.05, **p < 0.01, ***p<0.001 comparing conditions to control using Mann-Whitney tests. Scale bars represent 5μm in (A, E, F and H).

We first performed an unbiased screen of P180 interacting proteins by performing streptavidin pulldowns with GFP-biotin(GFPbio)-tagged P180 (**Figure S3B**) and subsequent mass spectrometry analysis using HEK293T lysates and adult rat brain extracts. In both cases, ribosomal proteins were amongst the top hits of interactors (**Figure 3C; Fig S3C**). Western blot analysis after streptavidin pulldowns further confirmed the interaction of ribosomal proteins with P180 (**Figure 3D**). Importantly, P180 also interacts with ribosomes in the axon since the split APEX assay with P180-AP and RpL10A-EX generated clear axonal biotinylation (**Figure 3E**).

Previous studies have shown that P180, which is enriched in ER sheets in non-neuronal cells, is important for ribosome/mRNA binding to the ER.^20,21,26^ We wondered whether P180, enriched in axonal ER tubules, could play a role in the interaction between the axonal ER and ribosomes, thus regulating axonal protein synthesis. We therefore investigated the effect of shRNA-mediated P180 KD^14^ on axonal translation. P180 KD caused a reduction in axonal protein synthesis, as measured by puromycilation (**Figure 3F and 3G**). To confirm that this reduction was due to impaired contacts between the axonal ER and ribosomes, we used the split APEX assay with AP-Sec61β and RpL10A-EX. This revealed a significant reduction in ER-ribosome interactions in the axon after P180 KD (**Figures 3H and I**).

Previous in vitro studies have suggested that P180 interacts with ribosomes through binding to mRNA.^20,27,28^ To determine if the interaction between P180 and ribosomal proteins is dependent on mRNA, we treated our pulldown samples with RNaseA/T1 to digest mRNA. This resulted in a significant decrease of ribosomal proteins bound to P180 (**Figure 3J and 3K**), suggesting that the interaction of P180 with ribosomes is stabilized by its binding to mRNA. These data are consistent with a model in which P180 regulates the targeting of mRNAs and ribosomes to the axonal ER membrane, thereby regulating the translation of a subset of mRNAs.

### ER – ribosome contacts require two binding domains in P180

It remains unclear which domains of P180 are required for ribosome/mRNA binding.^20,26,29^ We attempted to resolve this by studying the binding site(s) and function(s) on axonal ER – ribosome contacts using different P180 deletion constructs (**Figure 4A**). We first performed GFPbio pulldowns with full-length P180, P180ΔCoiled-coil (CC) and P180ΔRepeats. Neither the deletion of the CC domain nor the repeats domain reduced the interaction of P180 with ribosomes, compared to full-length P180 (**Figure 4B**). This may be explained by a possible dimerization of our expressed constructs with endogenous P180, as previously reported.^30^ Therefore, we expressed only the CC domain or only the repeats domain to see if one of these domains is the main binding site for ribosomes. However, pulldowns using only the CC, or the repeats domain revealed very little to no interaction with ribosomes (**Figure 4C**). To determine whether the association of P180 to the ER membrane or the lysine-rich domain is required for P180 – ribosome interactions, we expressed only the cytosolic domain containing both the repeats and CC domains. Surprisingly, this construct showed a strong interaction with ribosomes, similar to full-length P180 (**Figure 4D**). This shows that both the CC and repeats domains are necessary and sufficient to bind ribosomes.

**Figure 4.**
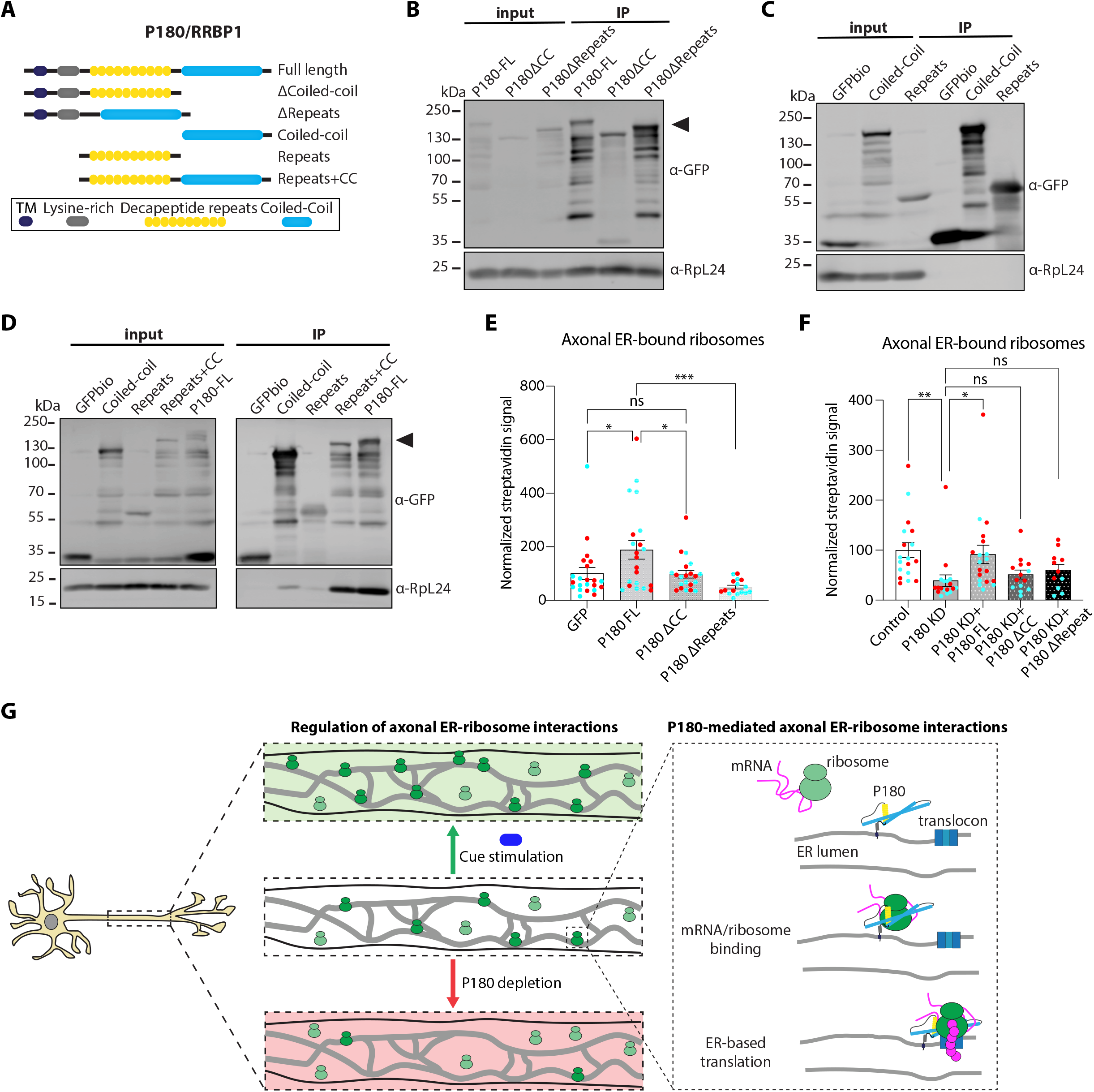
Axonal ER-ribosome interactions are regulated by specific P180 domains. (A) Schematic representation of P180 constructs used. (B, C, D) Representative Western blot analysis of GFP and endogenous RpL24 after GFPbio pulldowns with indicated constructs. (E-F) Quantification of axonal ER-bound ribosomes using split APEX assay in neurons expressing V5-AP-Sec61β and RpL10A-3xHA-EX, together with indicated P180 constructs in (E), or expression of the same constructs together with shRNAs targeting P180 in (F). (G) Schematic of our proposed model in which axonal ER-bound ribosomes and translation are regulated by P180 and extrinsic cues. Individual data points each represent a neuron and data is presented as mean values ± SEM in (E and F). ns = not significant, *p < 0.05, **p < 0.01, ***p<0.001 comparing conditions to each other using ordinary one-way ANOVA tests.

To determine whether P180, associated to the ER membrane but lacking its CC or repeat domain could still interfere with the recruitment of ribosomes to the axonal ER, we performed the split APEX assay with AP-Sec61β and RpL10A-EX. We found that while full-length P180 promotes axonal ER – ribosome contacts, neither P180ΔCC nor P180ΔRepeats was able to induce ER – ribosome interactions (**Figure 4E**). Importantly, we evaluated the requirement of these two different domains for proper axonal ER – ribosome interactions. We found that the drastic reduction in axonal ER – ribosome contacts caused by P180 KD, is rescued by full-length P180 but not P180 lacking the CC or the repeats domain (**Figure 4F**). Together, these results indicate that P180 is an axonal ribosome receptor with two essential binding sites required for proper ER – ribosome interactions.

## DISCUSSION

In this work, we find that axonal ER tubules play a key role in supporting local translation. The axonal ER binds ribosomes and is a site for local translation of at least a subset of mRNAs. The axonally enriched integral ER protein P180 acts as a ribosome/mRNA receptor that efficiently targets mRNA-bound ribosomes to the axonal ER for proper local translation. We propose a model in which P180 recognizes mRNA-containing ribosomes by two binding sites present in its repeats and CC domains. Axonal ER – ribosome interactions provide a subcellular platform for efficient protein synthesis, and these contacts are dynamically regulated by specific extracellular stimuli (**Figure 4G**).

Recent studies have shown that many mRNAs are present and being translated in the axon.^4,31,32^ Here, we show that axonal ER tubule disruption impairs overall local translation. Using four different super-resolution techniques, we consistently find that ribosomes form contacts with the axonal ER. Consistent with our findings, a recent study has visualized clusters of ribosomes close to the ER at axon branch points by Cryo-ET.^33^ The limited evidence of polysomes bound to axonal ER could be related to the possibility that ribosomes attached to the axonal ER consist of mainly monosomes, which are difficult to visualize by EM because of their small size. Monosomes were initially believed to be translationally inactive but were recently found to be actively involved in local translation, including translation of transmembrane, secretory and cytosolic proteins.^32^ Our super-resolution imaging data unfortunately does not provide the resolution needed to dissect between polysomes and monosomes. It thus remains unclear whether they are bound to different regions of the axonal ER; for instance, polysomes at axon branch points and monosomes along the axon shaft.

Importantly, we show that the axonal ER is a site for local translation. At present, our work is limited to the visualization of overall translation. We find that specific extrinsic stimuli regulate axonal ER-bound ribosomes. Cue-specific changes in the axonal proteome are known to occur.^31,34^ Interestingly, a previous study identified a specific upregulation of several transmembrane protein mRNAs in response to NT-3, but not BDNF or NGF stimulation.^34^ To fully understand the possible functions related to axonal ER-based translation, future studies are needed to identify which exact mRNAs are being translated at the axonal ER. This could reveal if the axonal ER is involved in the local synthesis of transmembrane/secretory proteins and possibly the translation of mRNAs coding for cytosolic proteins, as these can also be translated on the ER membrane in non-neuronal cells.^8^

Previous studies have shown a role for lysosomes and mitochondria in axonal mRNA localization and translation through RNA granule hitchhiking.^35–38^ The ER forms contacts with many different organelles, including mitochondria and lysosomes, but also with RNA granules.^39,40^ We observe that ER tubule disruption reduces local translation, which is caused by impaired ribosome distribution. It is possible that a mislocalization of mitochondria or lysosomes in the axon, due to impaired contacts with the axonal ER, affects mRNA localization and translation. However, mitochondria or lysosome transport into the axon are not affected by axonal ER disruption.^14^ We still have limited understanding of the possible role of organelle-organelle contacts in the distribution and dynamics of local translation in neurons. Our findings that the ER is involved in local mRNA translation opens the interesting possibility that ER-organelle contact sites are a location for mRNA exchange and translation between the ER and RNA granules or organelles on which RNA granules hitchhike.

We identify P180 as an integral ER membrane protein that facilitates axonal ER – ribosome interactions and influences local translation. P180 was initially identified as a ribosome receptor in non-neuronal cells^41^ but has also been suggested to target specific mRNAs to the ER, possibly via specific RNA-binding proteins (RBPs), thereby promoting ER-based translation.^27,28^ Indeed, we detected various RBPs in our P180 interactor screen **(Figure S3D**). Our findings of mRNA-dependent binding of ribosomes and P180-mediated recruitment of ribosomes to the ER are consistent with a model in which mRNA binding stabilizes ribosome interactions with the ER for local translation (**Figure 4G**).

We find that both the decapeptide repeats and coiled-coil domains are required for efficient ribosome binding, but deletion of either of these domains impairs ribosome targeting to the axonal ER. Based on a predicted structure of P180 using Alphafold^42^ (**Figure S4**) it is possible that a region spanning both domains is essential for maintaining a structural conformation that allows efficient binding and targeting of mRNAs and ribosomes to the ER. Since P180 is also known to bind and stabilize microtubules (MTs)^12,30^ it is possible that P180 provides a molecular location at which MT-transported RNA granules can dock for subsequent mRNA translation at the axonal ER. In addition, P180 could limit translocon mobility by stabilizing MTs, which has been shown to increase ribosome binding to the ER.^43^

Altogether, the results in this study indicate that the axonal ER plays a substantial role in regulating the local proteome. Although the functions of the axonal ER are poorly characterized^44^, the axonal ER has been shown to be important for axon development and synaptic function.^12,45^ This work opens the possibility that local ER-based translation plays a supportive role in these processes. Interestingly, both ER dysfunction, partly through mutations in ER proteins^46^, and dysregulation of local protein synthesis^47^ are reported in several neurodegenerative diseases. Our work provides a novel possible link between ER dysfunction and dysregulation of local translation in disease etiology.

## ACKNOWLEDGMENTS

This work was supported by the Netherlands Organization of Scientific Research (NWO Veni grant (VI.VENI.202.113) to M.K.; NWO Vidi grant (0.16.VIDI.189.019) to G.G.F.). and the European Research Council (ERC-StG 950617 to G.G.F.; ERC-CoG to C.C.H.; ER-CoG 819219 to L.C.K). J.M was supported by a EMBO long-term fellowship (ALTF 741-2020).

## AUTHOR CONTRIBUTIONS

M.K. designed and performed experiments, analyzed data, coordinated the study and wrote the manuscript. N.O performed experiments, analyzed data and edited the manuscript, H.N. performed STED imaging and analysis. D.J. performed SMLM imaging and J.M. performed TREx imaging under the supervision of L.C.K. R.S. performed mass spectrometry processing and analysis under supervision of M.A. L.C.K and C.C.H gave advice throughout the study and edited the manuscript. G.G.F. supervised and coordinated the study and wrote the manuscript.

## DECLARATION OF INTERESTS

C.C.H. is an employee of Genentech, a member of the Roche group. The authors declare no additional competing interests.

**Figure S1 (Related to Figure 1).**
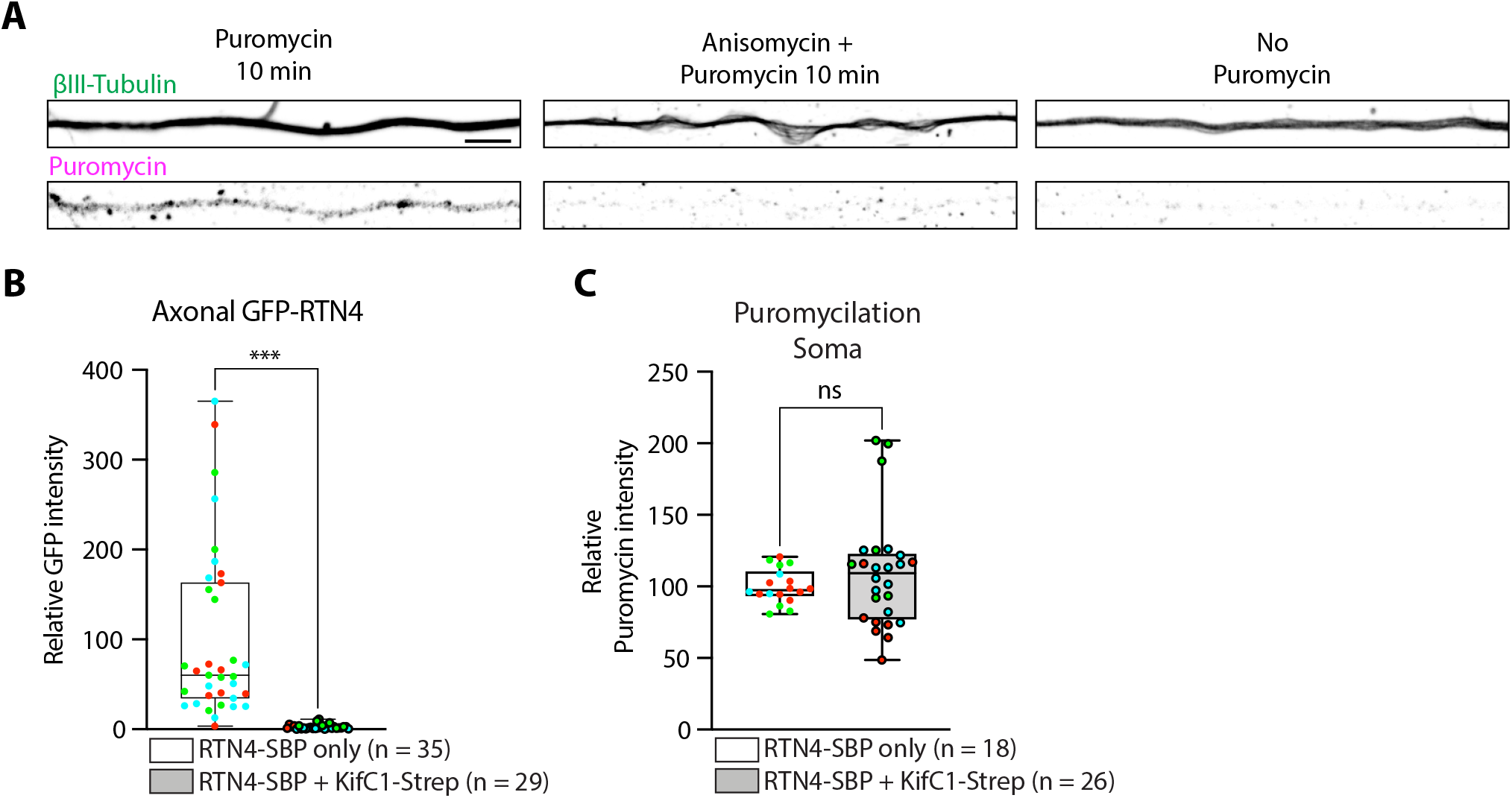
Axonal puromycilation specificity, efficient and selective ER removal from the axon which does not affect somatic puromycilation (A) Representative images of puromycilated peptides in distal axons from neurons treated with puromycin for 10 min (left), treated with puromycin for 10 min after anisomycin pre-treatment for 30 min (middle) or without puromycin (right). (B Quantification of RTN4-SBP-GFP levels in distal axons of DIV7 neurons expressing a fill plus RTN4-SBP-GFP in absence or presence of Strep-KifC1. (C) Quantification of puromycin intensity in neuronal soma of DIV7 neurons expressing a fill and RTN4-SBP-GFP in absence or presence of Strep-KifC1. Individual data points each represent a neuron and each color represents an independent experiment. Data are presented as mean values ± SEM in (B, C). ns = non-significant, ***p<0.001 comparing conditions to control using Mann-Whitney tests. Scale bars represent 5 μm (A).

**Figure S2 (Related to Figure 2).**
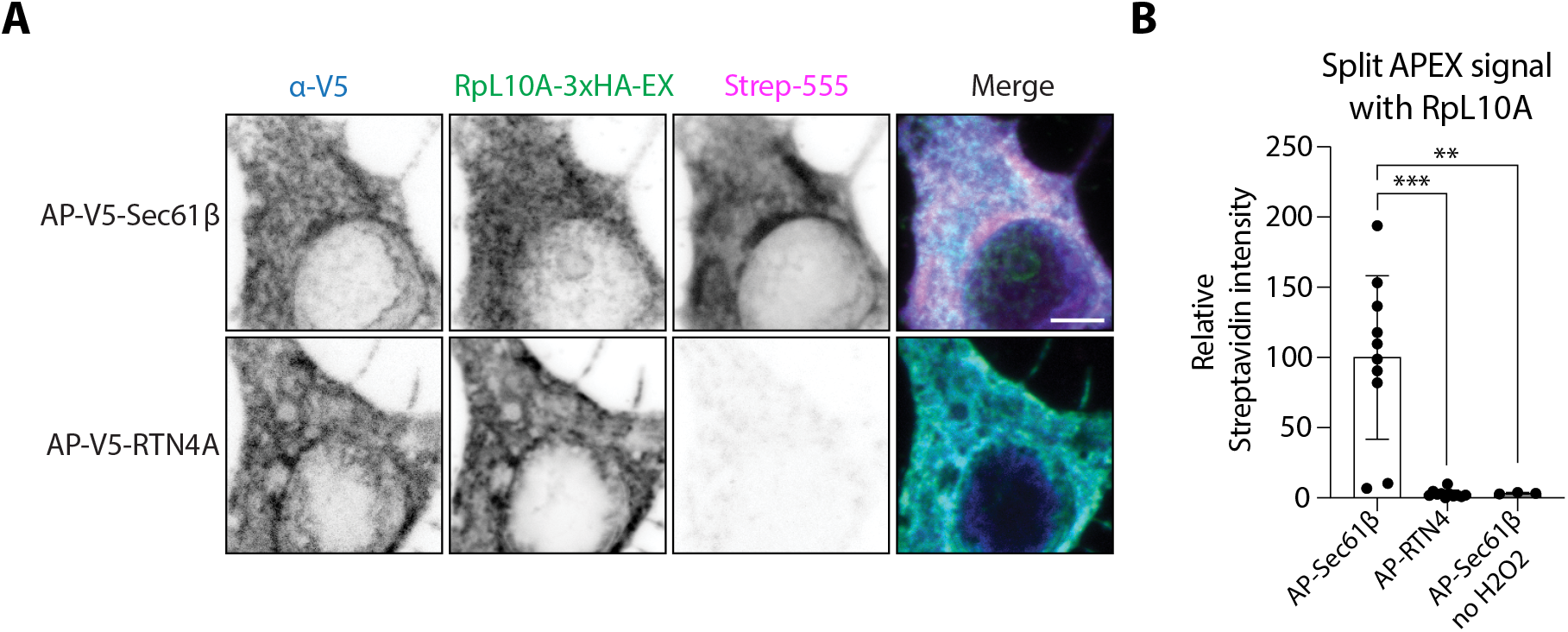
Validation of split APEX assay in neuronal soma (A) Representative images of split APEX assay in soma from neurons expressing Rpl10A-3xHA-EX and V5-AP-Sec61β (top), or V5-AP-RTN4A as a negative control (bottom). Expression of constructs are visualized with V5 and HA antibodies, and biotinylation is detected with conjugated Strep-555. Scale bars represent 5 μm. (B) Quantification of Strep signal in soma from neurons as in (A), and without H2O2 as a negative control for the biotinylation reaction. Individual data points each represent a neuron and data are presented as mean values ± SD in (B). **p < 0.01, ***p<0.001 comparing conditions to V5-AP-Sec61β using ordinary one-way ANOVA tests.

**Figure S3 (Related to Figure 3).**
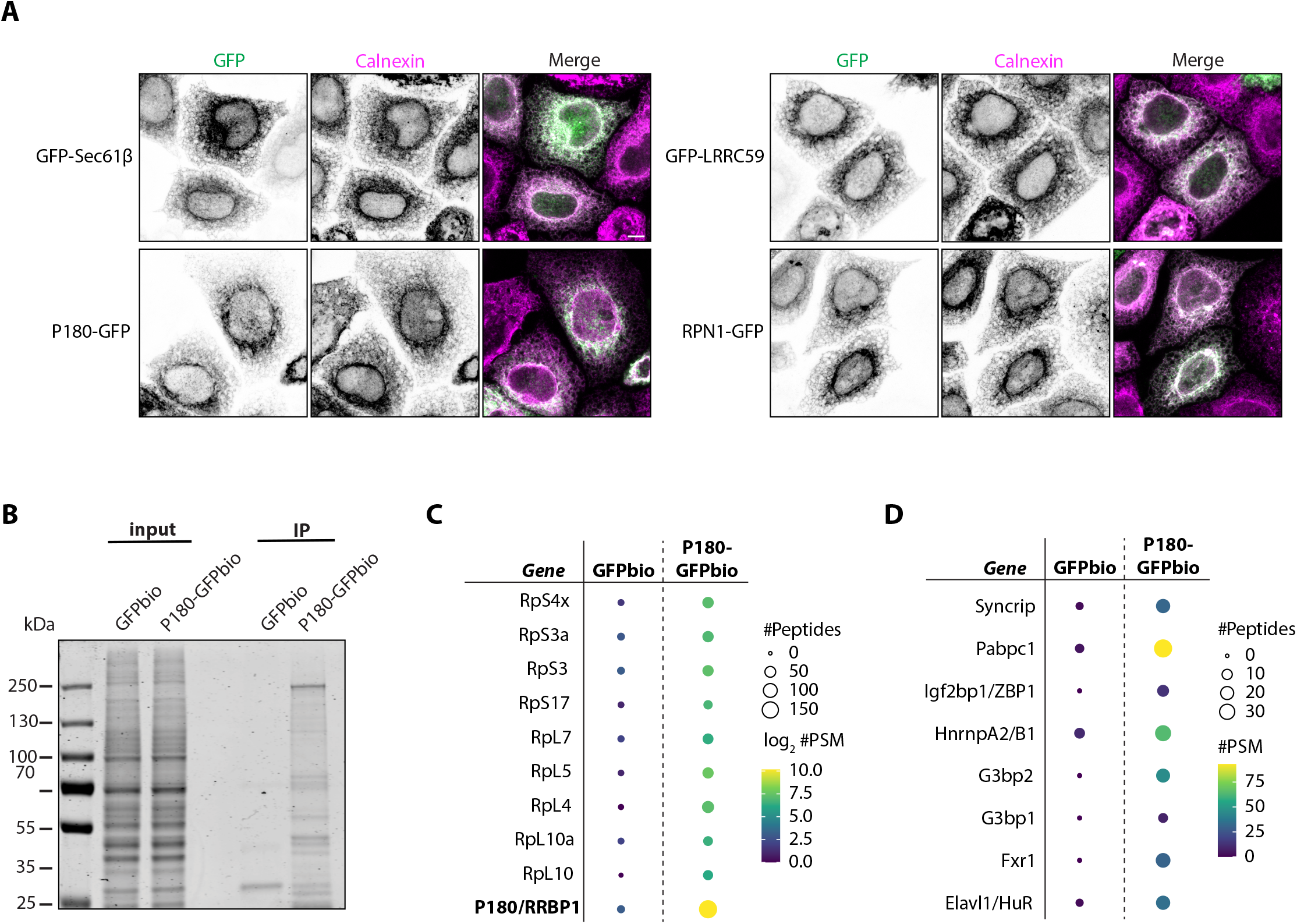
Correct ER localization of GFP-tagged constructs and ribosomal protein and RNA-binding protein interactions with P180 (A) Representative images of GFP-tagged ER protein constructs (as indicated) expressed in HEK293T cells and co-stained with an antibody against the ER marker Calnexin. All constructs show a correct ER localization. Scale bars represent 5 μm. (B) Silver-stained gel from input and pulldown samples of GFPbio and P180-GFPbio. (C) Scaled representation of ribosomal proteins identified with mass spectrometry after pulldown of GFPbio or P180-GFPbio from HEK293T cells. The size and color of each dot reflect the number of PSMs or peptides identified as indicated in the legend. (D) Scaled representation of RNA-binding proteins identified with mass spectrometry after pulldown of GFPbio or P180-GFPbio from adult rat brain extracts. The size and color of each dot reflect the number of PSMs or peptides identified as indicated in the legend.

**Figure S4 (Related to Figure 4).**
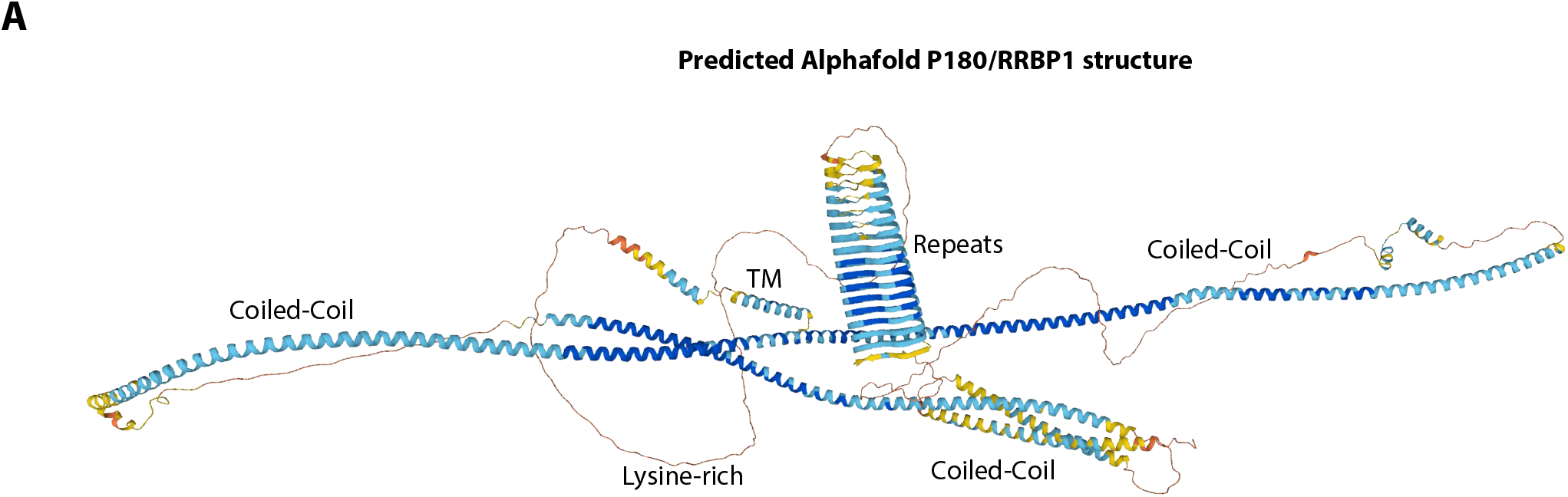
Predicted P180/RRBP1 structure (A) Alphafold2 predicted structure of P180/RRBP1 with each domain annotated.

## METHODS

### CONTACT FOR REAGENT AND RESOURCE SHARING

Further information and requests for resources and reagents should be directed to and will be fulfilled by the Lead Contact Ginny G. Farías.

### EXPERIMENTAL MODEL AND SUBJECT DETAILS

#### Animals

All experiments were approved by the DEC Dutch Animal Experiments Committee (Dier Experimenten Commissie), performed in line with institutional guidelines of University Utrecht, and conducted in agreement with Dutch law (Wet op de Dierproeven, 1996) and European regulations (Directive 2010/63/EU). Female pregnant Wistar rats were obtained from Janvier, and embryos (both genders) at E18 stage of development were used for primary cultures of hippocampal neurons. Brains from these female rats were used to obtain protein extracts. The animals, pregnant females and embryos have not been involved in previous procedures.

#### Primary neuronal cultures and transfection

Primary hippocampal neurons were prepared from embryonic day 18 rat brains from which the hippocampi were dissected, dissociated in trypsin for 15 min and 37°C and plated at a density of 100,000/well or 50,000/well (12-well plates) on coverslips coated with poly-L-lysine (37.5 μg/mL; Sigma) and laminin (1.25 μg/mL; Roche). Neurons were maintained in neurobasal medium (NB; Gibco) supplemented with 2x B27 (Gibco), 0.5 mM glutamine (Gibco), 15.6 μM glutamate (Sigma), and 1% penicillin/streptomycin (Gibco) and incubated under controlled temperature and CO_2_ conditions (37°C, 5% CO_2_).

Hippocampal neurons were transfected at day in vitro (DIV)3-4 using Lipofectamine 2000 (Invitrogen). Shortly, DNA (0.05-2 μg/well) was mixed with Lipofectamine 2000 (1.2 μL) in Opti-MEM (Gibco, 200 μL) and incubated for 20 min at room temperature. The mix was added to the neurons in NB without additives and incubated for 1 hour at 37°C in 5% CO2. Neurons were then washed with NB for 3 times and transferred back to their original medium at 37°C in 5% CO2 until fixation at DIV4 or DIV7.

#### HEK293T cell culture and transfection

Human embryonic kidney (HEK293T) cells (ATCC) were cultured in DMEM high glucose medium (Capricorn Scientific) supplemented with 10% fetal bovine serum (Gibco) and penicillin/streptomycin (Gibco). The cells were maintained at 37 °C in 5% CO2. HEK293T cells were plated into 10cm dishes (for mass spectrometry analysis) or in 60mm dishes (for pulldown-WB) and transfected using PEI MAX with different plasmids. Briefly, GFPbio constructs (varying amounts depending on the construct) together with a BirA plasmid (1:2.5 ratio) were mixed with PEI at a 1:2.5 ratio in Opti-MEM and incubated for 20 min at room temperature. Fresh DMEM medium with supplements was added to this mix and this was subsequently added to the cells which were placed back at 37 °C in 5% CO2. After 24-48 hours, cells were processed for biotin-GFP pulldowns or immunocytochemistry as described below.

### METHOD DETAILS

#### DNA and shRNA Constructs

Following vectors were used: pSuper^48^, pGW1-mCherry and pGW1-BFP^49^, RTN4A-GFP (a gift from Dr. Gia Voeltz, Addgene plasmid #61807), pEGFP(A206K)-N1 and pEGFP(A206K)-C1 (a gift from Dr. Jennifer Lippincott-Schwartz), GFP-Sec61β (a gift from Dr. Tom Rapoport, Addgene # 15108), RpL10A-tagRFP (a gift from dr. Robert Singer, Addgene #74172) and TOM20-V5-FKBP-AP (a gift from Alice Ting, Addgene#120914). HA-KifC1-MD-Strep, GFP-SBP-RTN4A, P180-ΔCoiled-coil, P180-Δrepeats-GFP, P180-Coiled-coil and P180-Repeats were previously described.^12^ SplitAP-V5-C1 and 3xHA-split-EX-N1 were previously described.^14^

The plasmids generated in this study include:

For GFP-LRRC59, LRRC59 sequence was PCR amplified from a cDNA library generated from a rat INS-1 cell line and inserted in pEGFP(A206K)-C1 vector between BglII and BamHI sites using HiFi DNA assembly (New England Biosciences). A flexible linker was added before LRRC59 by addition to the cloning primers. Primers used to generate GFP-LRRC59 constructs were as follows:

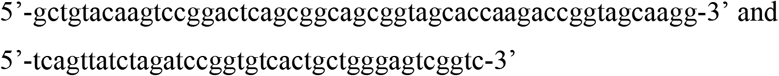

For RPN1-GFP, RPN1 sequence was PCR amplified from a rat INS-1 cDNA library and inserted in pEGFP(A206K)-N1 vector between HindIII and AgeI sites by HiFi DNA assembly. Kozak sequence was generated by addition to the cloning primers and was introduced before RPN1 sequence. A flexible linker was inserted between RPN1 and GFP by addition to the cloning primers. The primers used to generate RPN1-GFP were:

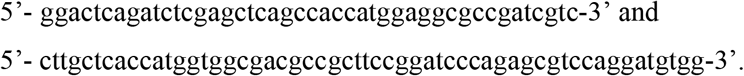

For KTN1-mNG, human KTN1 was PCR amplified from Clone I.M.A.G.E: 40125683 and inserted into pmNeonGreen-N1, between BamHI and XhoI restriction sites by HiFi DNA assembly. To generate KTN1a, the longest KTN1 isoform, DNA sequences missing from the IMAGE clone were subsequently added between aa-831 and aa-855, and between aa aa-1229 and aa-1258 by PCR.

For P180-Repeats+Coiled-coil construct, the DNA sequence between aa194 and aa1540 was amplified from V5-GFP-P180 (Addgene #92150), and this fragment was cloned into pEGFP(A206K)-N1 between BamHI and XhoI sites by HiFi DNA assembly. A kozak sequence, start codon and flexible linker were inserted by addition to cloning primers. The primers used to generate this construct are:

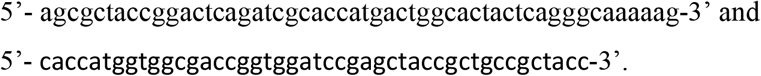

For GFP-AviTag versions of all P180 constructs, a GFP-AviTAG fragment was PCR amplified from GFP-AviTag-N1 and inserted into the GFP-tagged constructs between BamHI and BsrGI sites (which removes the existing GFP) by HiFi DNA assembly.

For Split APEX assay, Split-AP-V5-Sec61β, RpL10A-3xHA-Split-EX and P180-V5-Split-AP were generated.

For Split-AP-V5-Sec61β, Sec61β sequence was PCR amplified from GFP-Sec61β and inserted in Split-AP-V5-C1 vector between BglII and EcoRI. A flexible linker was introduced before Sec61β. The primers used to generate Split-AP-V5-Sec61β were:

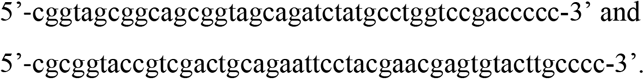

For RpL10A-3xHA-Split-EX, RpL10A sequence was PCR amplified from RpL10A-TagRFP and inserted in 3xHA-Split-EX-N1 vector between AgeI and HindIII. A flexible linker was introduced after RpL10A. The primers used to generate RpL10A-3xHA-Split-EX were:

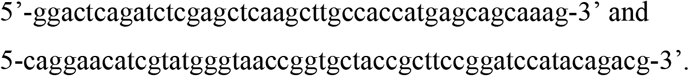

For P180-V5-Split-AP, V5-Split-AP was PCR amplified from TOM20-V5-AP and inserted in P180-FL-GFP vector between BsrGI and BamHI sites (which removes existing GFP). A flexible linker was added after P180 sequence. The primers used to generate P180-V5-Split-AP were:

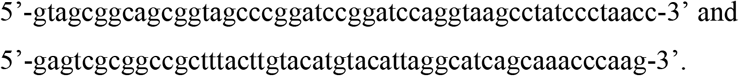

For Split-AP-V5-RTN4, RTN4 sequence was PCR amplified from GFP-SBP-RTN4 and inserted into Split-AP-V5-C1 vector between BglII and EcoRI restriction sites. A flexible linker was introduced before RTN4. The primers used to generate Split-AP-V5-RTN4 were:

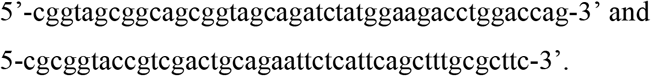

The following sequences for rat-shRNAs were used in this study: RTN4-shRNA (5’-gtccagatttctctaatta-3’), DP1-shRNA (5’-gacatataaagttccagaa-3’) validated in Farias et al., 2019^12^; P180-shRNAs (5’-tcagtgcaattgtctgtat-3’ and 5’-taaaccaaccaacacagcg-3’) validated in Özkan et al., 2021.^14^

#### Antibodies and reagents

The following primary antibodies were used in this study: mouse anti-Puromycin (Kerafast Cat# EQ0001, RRID:AB_2620162, 1:3000), rabbit anti-RpS12 (Proteintech Cat#16490-1-AP, RRID:AB_2146233, 1:200 for IC, 1:1000 for WB), rabbit anti-RpL24 (Proteintech Cat# 17082-1-AP, RRID:AB_2181728, 1:200 for IC, 1:1000 for WB), mouse anti-V5 (Thermo Fisher Scientific Cat# R960-25, RRID:AB_2556564, 1:1000 for IC), rat anti-HA (Roche Cat# 11867423001, RRID:AB_390918, 1:1000 for IC), mouse-anti-GFP (Thermo Fisher Scientific Cat#A-11120; RRID:AB_221568, 1:250-500 for IC), rabbit-anti-GFP (Abcam Cat#ab289; RRID:AB_303395, 1:10.000 for WB).

The following secondary antibodies were used in this study: Alexa-Fluor555 conjugated Strep (Thermo Fisher Scientific Cat# s21381, RRID:AB_2307336, 1:2000), Alexa-Fluor-568 conjugated Strep (Thermo Fisher Scientific Cat# S-11226, RRID:AB_2315774, 1:1000), goat anti-Mouse IgG (H+L) Highly Cross-Adsorbed Alexa Fluor 488 (Thermo Fisher Scientific Cat# A-11029, RRID:AB_2534088, 1:1000 for IC, 1:200 for TREx), goat anti-mouse Alexa568 (Thermo Fisher; Cat#A-11031, RRID:AB_144696, 1:1000), goat anti-mouse Alexa405 (Molecular Probes, Cat# A31553, RRID:AB_221604, 1:500), goat anti-rat Alexa 488 (Thermo Fisher Scientific Cat# A-11006, RRID:AB_2534074, 1:1000). Anti-rabbit IRDye 800CW (Li-Cor, Cat#926-32211, RRID:AB_621848), Atto488 conjugated Fluotag-X4® GFP nanobody (NanoTag Biotechnologies, Cat# N0304-At488-L, 1:250), goat anti-rabbit IgG (H+L) Highly Cross-Adsorbed CF®594 (Sigma-Aldrich, Cat#SAB4600110, 1:1000), goat anti-mouse IgG Abberior STAR 635P(Abberior Cat# ST635P-1001-500UG, RRID:AB_2893232, 1:500), goat anti-rabbit Alexa 647 (Thermo Fisher Scientific Cat# A-21245, RRID:AB_2535813, 1:500), goat anti-mouse CF680 (Biotium Cat# 20065, RRID: AB_10557108, 1:500), anti-rabbit ATTO 647N (Sigma-Aldrich, Cat#40839, RRID:AB_1137669, 1:200).

Other reagents used in this study were: Puromycin dihydrochloride (Sigma-Aldrich, Cat# P8833), Anisomycin (Sigma-Aldrich, Cat#9789), recombinant human Neurotrophin-3 (NT-3) protein (50ng/ml, Alomone labs, Cat#N-260), recombinant human BDNF protein (50ng/ml, Alomone labs, Cat#B-250), recombinant rat beta-NGF Protein (50ng/ml, R&D systems, Cat# 556-NG). NeutrAvidin (Thermo Fisher Scientific, Cat# 31000), Heme (Sigma-Aldrich, Cat# 51280); biotin-phenol (Iris Biotech, Cat#LS.3500); H_2_O_2_ (Sigma-Aldrich, Cat#H1009), Lipofectamine 2000 (ThermoFisher Scientific, Cat#11668019), Polyethylenimine (PEI MAX; Polysciences, Cat#24765), Fluoromount-G Mounting Medium (ThermoFisher Scientific, Cat#00-4958-02), Dynabeads Streptavidin (Thermo Fisher, Cat#11205D), Pierce Streptavidin Magnetic beads (Thermo Fisher, Cat#88816), RNase A (Thermo Fisher, Cat#EN0531), RNaseT1 (Thermo Fisher, Cat#EN0541), Trolox (Sigma-Aldrich, Cat#238831), Sodium L-Ascorbate (Sigma-Aldrich, Cat#A4034), Sodium Azide (Sigma-Aldrich, Cat#S2002), Cysteamine (MEA) (Sigma, Cat#30070), Glucose-oxidase (Sigma, Cat#G2133), Catalase (Sigma, Cat#C40), Acryloyl-X-SE (AcX, Thermo Fisher Scientific, Cat# A20770), Acrylamide (40%, Sigma, Cat#A4058-100ML), N,N′-Methylenebisacrylamide (bisacrylamide, Sigma, Cat#M1533-25ML), Sodium acrylate (Sigma, Cat# 408220-25g), TEMED (Bio-Rad, Cat#1610800), APS (Sigma, Cat#A3678), anhydrous-DMSO (Thermo-Fisher Scientific, Cat# D12345), Guanidine HCl (Sigma, Cat#G3272), Triton-X-100 (Sigma, Cat#93433), Proteinase K (Thermo Fisher Scientific, Cat#EO0492) and 0.1% (w/v) poly-L-lysine (Sigma, Cat#P8920-100mL).

### Puromycilation

In order to label newly synthesized proteins, we briefly incubated neurons with 10μM puromycin for 10 minutes. After 10 minutes, unincorporated puromycin was washed out using two washes with NB medium and neurons were fixed immediately and processed for immunostaining as described below.

#### Immunofluorescence staining and confocal imaging

Neurons were fixed with pre-warmed 4% paraformaldehyde plus 4% sucrose in PBS for 10-20 min at RT and washed with PBS supplemented with calcium and magnesium (PBS-CM) three times. Fixed cells were subsequently permeabilized with 0.2% Triton X-100 in PBS-CM for 15 min at RT, washed 3 times and were then incubated with blocking buffer (0.2% porcine gelatin in PBS-CM) for 1 hour at RT. Next, neurons were incubated with primary (1 hour at RT or O/N at 4°C) and secondary antibodies (1h at RT) at specified concentrations in blocking buffer. Coverslips were mounted in Fluoromount-G Mounting Medium and imaged by using a confocal laser-scanning microscope (LSM700, with Zen imaging software (Zeiss) version 8.1.7.484) equipped with Plan-Apochromat ×63 NA 1.40 oil DIC and EC Plan-Neofluar x40 NA1.30 Oil DIC objectives.

#### Stimulated Emission Depletion (STED) microscopy

Imaging was performed with a Leica TCS SP8 STED 3x microscope using an HC PL APO 100x/NA 1.4 oil immersion STED WHITE objective. The 488, 561 and 633 nm wavelengths of pulsed white laser (80MHz) were used to excite Atto488, CF®594 and Star635P, respectively. To obtain gSTED images, Atto488 was depleted with the 592 nm continuous wave depletion laser, CF®594 and Star635P were depleted with the 775 nm pulsed depletion laser. An internal Leica HyD hybrid detector with a time gate of 0.3 to 6 ns was used. Images were acquired as Z-stack and maximum intensity projections were obtained for image display and analysis.

#### Ten-fold Robust Expansion (TREx) microscopy

Ten-fold robust expansion microscopy was performed as previously described^18^ with slight alterations. Briefly, neurons were fixed and immunostained as described above (using a two-fold higher concentration for primary antibodies) after which they were anchored with 100ug/ml acryloyl-X SE in PBS overnight at RT. After rinsing in PBS, coverslips containing the cells were inverted cell-side down onto pre-cooled gelation chambers (using silicon rings (13mm diameter, 120μL volume, Sigma, Cat# GBL664107)), containing the gelation solution (1.08 M sodium acrylate, 14.4% (v/v) acrylamide, 0.009% (v/v) bis-acrylamide, 1.5% TEMED, 0.15% (w/v) APS in PBS) on ice. Gelation occurred at 37°C for 30 min. Samples were subsequently rinsed in PBS and submerged in digestion buffer (0.5% Triton-X-100, 0.8M Guanidine-HCl, 9U/mL Proteinase K in TAE buffer (40mM Tris, 20mM acetic acid, 1mM EDTA) and digested for 4h at 37°C, followed by brief rinsing with MilliQ (MQ) water.

Samples were cut into smaller gel pieces to allow for faster expansion and transferred into 15cm cell culture dishes filled with MQ to promote physical sample expansion through osmosis (fresh MQ was exchanged three times over a course of 2 days to allow for complete sample expansion with an estimated expansion factor of ∼9 calculated using the overall increase in size of the gel measured before mounting and imaging the sample, compared to the initial gel size after gelation). Samples were subsequently cut with a razor and mounted on 3D printed sample holders (printed with PLA using a Prusa mini printer and a model generated in Fusion360, see .obj file) onto a poly-L-lysine coated, and plasma-cleaned, rectangular coverslip.

Imaging was performed on a Leica SP8 3X STED microscope using a 63x (HC PL APO CS2, NA1.20) water objective using confocal setup. Detailed acquisition settings were the following: bidirectional scanning, 400Hz speed, zoom factor 7, 1AU pinhole, twofold frame averaging, 30% laser intensity (at 488nm, 8-fold line accumulation, detection using a HyD at 495nm - 570nm and gating set to 0.3ns – 6ns) and 30% laser intensity (at 633nm, three-fold line accumulation, detection using a HyD at 644nm - 736nm and gating set to 0.3ns – 6ns). Images were acquired using the sequential image settings as a z-stack over a physical distance of 0.92nm using three slices to capture the axon.

#### Dual-color Single Molecule Localization Microscopy (SMLM)

Dual-color SMLMwas performed as previously described.^19^ Briefly, we used a method similar to spectral demixing, to classify two spectrally very close far-red fluorophores enabling high-resolution stochastic optical reconstruction microscopy (STORM) imaging of two channels. Neurons were fixed and immunostained as described above using goat anti-rabbit AlexaFluor (AF) 647 and goat anti-mouse CF680 secondaries. Samples were mounted in imaging buffer (100mM MEA, 5% w/v glucose, 700 μg/ml glucose oxidase, 40 μg/ml catalase in 50mM Tris pH 8.0) in closed off cavity slides (Sigma, BR475505) to prevent oxygen from entering the sample during imaging. Imaging was performed on a Nikon TI-E microscope equipped with a TIRF APO x100 NA 1.49 oil objective lens. A 638 nm laser (MM, 500mW, Omicron) together with a laser clean-up filter (LL01-638, Semrock) and excitation dichroic (FF649-Di01, Semrock) was used to excite the sample. The collected emission was relayed through an Optosplit III module (Cairn Research), fitted with emission dichroic (FF660-Di02) to split the emission in a short channel and a long channel on a EMCCD (iXon 897 – Andor). Samples were imaged with laser at an TIRF angle, for 16000 frames with an exposure of 10 ms.

Acquisitions were analyzed using the custom ImageJ plugin DoM (Detection of Molecules, https://github.com/ekatrukha/DoM_Utrecht), and reconstructions were generated by plotting the resulting localizations. Further analysis was performed using PFC (Probability-based Fluorophore Classification).^19^ Using the short channel for classification with a generalized likelihood ratio test (GLRT) and the long channel for localization, fluorophores AF647 and CF680 are separated. The result is two high resolution reconstructions of our proteins of interest. For Figure 2K, neurons were treated with a high concentration (200μM) of puromycin for 45 minutes before fixation and processing as described above.

#### Streptavidin/SBP heterodimerization system assay

Controlled coupling of ER tubules to MT-driven motor proteins has been previously described.^12^ Briefly, neurons were transfected at DIV4 with only GFP-SBP-RTN4A as a control or GFP-SBP-RTN4A plus HA-KifC1-MD-Strep to pull axonal ER tubules to soma. NeutrAvidin (0.3 mg/ml) was added to the neurons after 1 h of transfection to prevent Strep-SBP uncoupling.

#### Split APEX assay

Split APEX assay was performed as described previously.^14^ Briefly, neurons were transfected at DIV4 with RpL10A-HA-EX and AP-V5-Sec61β, V5-AP-RTN4A or P180-V5-AP constructs. At DIV7, neurons first were incubated with heme (6 μM for 60 min at 37°C/ 5% CO2 and subsequently washed once with NB and incubated with biotin-phenol (500 μM) in NB with supplements for 30 min at 37°C/ 5% CO2. For cue stimulation experiments, cues were added together with biotin phenol for 30 min at 50ng/ml. Then, H2O2 to a final concentration of 1 mM was introduced to the medium for 1-2 min to initiate proximity labelling, after which the reaction was stopped by removing the medium and washing the cells once with quenching buffer (5 mM Trolox and 10 mM sodium ascorbate in HBSS) containing 10 mM sodium azide and twice with quenching buffer without sodium azide for 3–5 min each at 37°C/ 5% CO2. Neurons were subsequently fixed and immunostained as described above.

#### Biotin-GFP pulldown assays

Dynabeads-M280-Streptavidin or Pierce Streptavidin magnetic beads were first blocked with a blocking buffer (20 mM Tris HCl pH7.5, 150 mM KCl and 0.2 ug/ul chicken egg albumin) for 1 hour by rotating at RT at 16 rpm and were then washed with wash buffer (20 mM Tris HCl pH7.5, 150 mM NaCl, 0.1% Triton-X 100 and 5 mM MgCl2) for 3 times using a magnetic rack.

Cells were collected by first washing them with ice-cold 1x PBS supplemented with 0.5x protease inhibitor cocktail (Roche) and then collecting them with a cell scraper in PBS/0.5x protease inhibitor in an eppendorf tube. Cells were pelleted by centrifugation at 3000g for 5 min at 4 °C. Cell pellets were resuspended in lysis buffer (20 mM Tris HCl pH7.5, 100 mM NaCl, 1% Triton-X 100, 5 mM MgCl2 and 1x protease inhibitor cocktail and then subsequently incubated at 4 °C by rotating at 16 rpm for 15-30 min. Lysed cells were cleared by centrifugation at 4 °C, 16100g for 20 min. 5% of the total lysate was kept as input sample and the remaining lysate was added to streptavidin beads and incubated at 4°C by rotating at 16 rpm for 1 hour. After this, beads were washed 4 times with 400ul wash buffer on a magnetic rack. For RNAse A/T1 treament, RNAse A (4ng/μl) and RNaseT1 (250U/ml) were added to wash buffer and beads were wash 4 times on a magnetic rack. Finally, beads were dissolved in 2x Leammli sample buffer with DTT and boiled at 95 °C for 10 min and the sample was then separated from the beads on a magnetic rack and transferred to a new tube. Samples were stored at –20 °C until processing for mass spectrometry or Western blot. For pulldowns using rat brain lysates, beads incubated with HEK293T lysates as above were washed twice with low salt buffer (100mM KCl, 0.1% TritonX-100, 20mM Tris, pH 7.6), twice with high salt buffer (500mM KCl, 0.1% TritonX-100, 20mM Tris, pH 7.6) and twice again with low salt buffer to remove binding proteins from HEK293T cells. Beads were then incubated with whole rat brain extract for 1h at 4°C and subsequently washed 5 times using normal wash buffer and sample was then collected from the beads as above.

#### Western blot analysis

Samples were loaded on a home-made 9 or 10% Bis-Acrylamide (Bio-Rad) gel and the gel was subsequently transferred by wet transfer to a PVDF membrane (Bio-Rad) for 90 min at 100V. The blots were blocked in blocking buffer (5% skimmed milk in TBS-T) for 1 h at room temperature. Blots were then incubated with primary antibodies diluted in blocking buffer at desired concentrations overnight at 4 °C on a rotator. Blots were washed with TBS-T 3 times for 5 min each on a shaker and incubated with secondary antibody (Li-Cor) in blocking buffer at desired concentrations for 1 h at room temperature in the dark. Finally, blots were washed 3 times with TBS-T for 5 min each and twice briefly in TBS before developing on an Odyssey CLx imaging system (Li-Cor) with Image Studio version 5.2 software. Protein levels were analyzed and normalized by importing images from Western blot detection into Fiji/ImageJ. SDS-PAGE silver stain was performed using a Pierce Silver Stain Kit (Thermo Fisher; Cat#24612).

#### Mass Spectrometry sample preparation and analysis

For mass spectrometry analysis, pulldown samples collected as described above were loaded on a 4-12% gradient Criterion XT Bis-Tris precast gel (Bio-Rad). The gel was fixed in 40% methanol and 10% acetic acid and subsequently stained for 1h using colloidal coomassie dye G-250 (Gel Code Blue Stain Reagent, Thermo Fisher). Each lane from the gel was cut in 3 gel pieces and placed in 0.5-mL tubes. Gel pieces were washed with water, followed by 15 min dehydration in acetonitrile. Proteins were reduced (10mM DTT for 1h at 56°C), dehydrated and alkylated (55 mM iodoacetamide for 1h in the dark). After two rounds of dehydration, digestion was performed by adding trypsin (Promega; 20μl of 0.1 mg/ml in 50 mM ammonium bicarbonate) and incubating overnight at 37°C. Peptides were extracted with acetonitrile, dried down and reconstituted in 10% formic acid.

All samples were analyzed on an Orbitrap Q-Exactive mass spectrometer (Thermo Fisher) coupled to an Agilent 1290 Infinity LC (Agilent Technologies). Peptides were loaded onto a trap column (Reprosil C18, 3μm, 2cm x 100μm) with solvent A (0.1% formic acid in water) at a maximum pressure of 800 bar and chromotographically separated over the analytical column (Zorbax SB-C18, 1.8μm, 40cm x50μm; Agilent) using 90 min linear gradient from 7-30% solvent B (0.1% formic acid in acetonitrile) at a flow rate of 150 nL/min. The mass spectrometer was used in a data-dependent mode, which automatically switched between MS and MS/MS. After a survey scan from 350-1500 m/z the 10 most abundant peptides were subjected to HCD fragmentation.

For data analysis, raw files were processed using Proteome Discoverer 1.4 (v1.4.14, Thermo Fisher). Database searches were performed using Mascot as a search engine (v2.5.1, Matrix Science) on the Human and Rat Uniprot databases. Carbamidomethylation of cysteines and oxidation of methionine were set as fixed and variable modifications respectively. Trypsin was set as cleavage specificity, allowing a maximum of 2 missed cleavages. Data filtering was performed using percolator, resulting in 1% false discovery rat. Additional filters were search engine rank 1 and mascot ion score >20. To infer protein abundance of each protein pulled down with the bait protein, we relied on total numbers of peptide spectrum matches (PSM). Dot plots of PSM and peptide numbers for selected proteins were generated using a custom-made script in R (R-project).

#### Image analysis and quantification

##### Puromycilation and ribosomal protein intensity

To analyze puromycilated peptide and ribosomal protein intensities, samples were imaged with the same settings for laser power, exposure and gain for all conditions. Distal axonal segments were selected for imaging by moving along the axon using a cell fill until the axon tip was reached. Next, an image of an axonal segment in this distal region that did not have many crossing neurites from other, untransfected neurons, was selected for imaging. Fiji/ImageJ was used to quantify the background corrected intensity. Average z-projections from acquired images were generated and segmented lines were manually drawn along axon segments (1-4 segments per neuron). Mean background corrected intensities from 16-bit images were measured.

##### Polarity index of ER proteins

Quantification of polarized distribution of proteins/organelles in neurons, polarity index (PI) has been previously described (Kapitein et al., 2010). Shortly, Fiji/ImageJ was used to draw segmented lines along an axonal region of ∼200 μm after the axon initial segment, and three dendrites per neuron. Mean intensities in axon and dendrites were measured. Following formula was applied PI = (Id-Ia)/(Id+Ia): where Id is the average mean intensity of the three dendrites and Ia is the mean intensity of axon. Non-polarized distribution represented by PI = 0 where Id = Ia, PI<0 indicates axonal distribution and PI>0 indicates dendritic distribution.

##### Split APEX streptavidin intensity quantification

For streptavidin signal intensity analysis, somata or axonal segments were imaged with the same settings for laser power, exposure and gain for all conditions. Axonal segments were imaged as. Fiji/ImageJ was used to quantify the background corrected intensity of signals. Average z-projections of images were generated and segmented lines were manually drawn along the axon. Mean intensities from 16-bit images were measured. The intensities for the strep signal were normalized to control conditions and per experiment. For comparison of Sec61β and RTN4 split APEX, the streptavidin intensities were additionally correct for V5 and HA intensities.

##### Quantification of ribosomes bound to ER with superresolution microscopy

To quantify the portion of ribosomes in contact with the axonal ER from STED, TREx and dual-color SMLM images, axonal segments were first straightened in Fiji/ImageJ. The mean intensity of the ribosomal channel was measured first. Then, masks from the Sec61β-GFP (ER) channel were generated by thresholding this channel. An outline of this mask was created and the intensity of the ribosomal channel within this mask was measured. For ‘ER enlarged’ the mask was enlarged with 5nm and mean ribosomal protein intensity was measured. The proportion of ribosomal protein intensity within the mask from the total amount was then calculated and plotted.

### Statistical analysis

Data processing and statistical analysis were performed using Microsoft Excel and GraphPad Prism. Unpaired t-tests, Mann-Whitney tests, ordinary one-way ANOVA tests followed by Dunnett’s multiple comparisons tests were performed for statistical analysis as indicated in figure legends.

